# Phylodynamic analysis reveals asymmetric and accelerating global spread of dengue virus

**DOI:** 10.64898/2026.09.28.754967

**Authors:** Verity Hill, Gytis Dudas, Xiang Ji, Ralph Huits, Davidson H. Hamer, Har-Kishen Joshi, Abigail J. Porzucek, Rafael Lopes, Hailey Robertson, Charles F. Whittaker, Moritz U.G. Kraemer, Michael A. Johansson, Nuno R. Faria, Oliver Brady, Simon Dellicour, Marc A. Suchard, Philippe Lemey, Colin J. Carlson, Guy Baele, Nathan D. Grubaugh

## Abstract

Dengue virus (DENV) continues to expand globally, with increasing incidence in both endemic and non-endemic countries. Leveraging a large global dataset of DENV genomes starting from the 1940s, we reconstructed global and regional patterns of DENV spread using phylogeographic and phylodynamic methods. Our analyses reveal that the current global diversity of urban DENV originated in Asia in the first half of the 20th century. During the more than 100 years since, we find that the highest density of between-country spread occurs within Asia and the region continues to be a global source, seeding transmission in Africa and the Americas. Global spread is, however, not balanced. Even though there is more travel to South America, we show that DENV lineages primarily enter North America (mostly from South Asia) through a link defined by seasonal synchrony across the Northern Hemisphere. Along with a substantial increase in global travel starting ∼2010, the rate of spread to the Americas is increasing, establishing new lineages associated with large recent outbreaks. DENV lineages do not frequently spread back to Asia from the Americas or Africa, and we posit this is due to higher genetic diversity in Asia, leading to stiffer competition for invading lineages. Within the Americas, the network of DENV dispersal initiates in North America, primarily the Caribbean, and then is maintained by major hubs in South America. Overall, we find that travel volume, rather than climate factors, is mostly responsible for long-term patterns of DENV spread globally and within the Americas. Our results highlight key nodes in the global dynamics of DENV lineages, and suggest spread will continue to accelerate in a more connected world.

## Introduction

The global expansion and increasing burden of dengue virus (DENV) is one of the most dramatic of the past century, but what drives regional prevalence and the underlying global transmission network remains poorly characterized. It is estimated that the four DENV serotypes originated ∼1000 years ago (*1*, *2*), each establishing urban transmission cycles in Southeast and South Asia by the late 1800s to early 1900s (*2*–*5*) with subsequent spread to the Pacific, Africa, and the Americas via human movement.

Phylogeographic studies have characterized some of the dispersal of DENV, including the establishment of Asian DENV genotypes in the Americas beginning in the 1970s-80s (*6*) and the recent global expansion of DENV-2 genotype II (“cosmopolitan”) (*7*–*11*). There are also several examples linking DENV lineage introductions to outbreaks, especially in the Americas (*12*–*18*). Yet, the global DENV lineage dynamics are complex due to fluctuations in transmission intensity (mostly due to climate) and heterogeneous dispersal networks (mostly due to human behaviors) (*19*, *20*). Furthermore, with the range of the *Aedes* spp. mosquito vectors increasing due to human-caused climate change and urbanization (*21*), there is potential for consequential new introductions to regions that were previously unsuitable for transmission. Despite the importance of these global dynamics, we do not fully understand the rates and drivers of spread across all four DENV serotypes at both global and regional scales.

Here, we leverage recent increases in global sequencing capacity, methodological innovation in phylodynamics (the co-analysis of genomic, epidemiological and immunological factors (*22*)), and newly sequenced DENV genomes from the mid-20th century (*23*) to build a unified analytical framework to study global DENV lineage dynamics. Our work reveals the directionality and intensity of DENV dispersal, permitting better targeting of surveillance resources and prediction of future spread.

## Results

### Asia is an important source for global DENV spread

To explore the dynamics of global DENV spread, including the most important intercontinental connections, we performed phylogeographic analyses on a cross-continental scale for each serotype (DENV-1-4; **Figure 1**). The timing and location of basal nodes can be affected by sampling bias (*24*), which can be improved even with a small number of genomes from under-represented locations or years (*25*). We therefore included 127 new DENV genomes from 1944 to 2016 that we sequenced from the Yale Arboviral Research Unit archives, representing all defined genotypes that have caused human outbreak (*23*). To further mitigate this bias, we downsampled the global genomic dataset to balance the sequences between South America and Southeast Asia (as major endemic regions; **Methods**).

**Figure 1:**
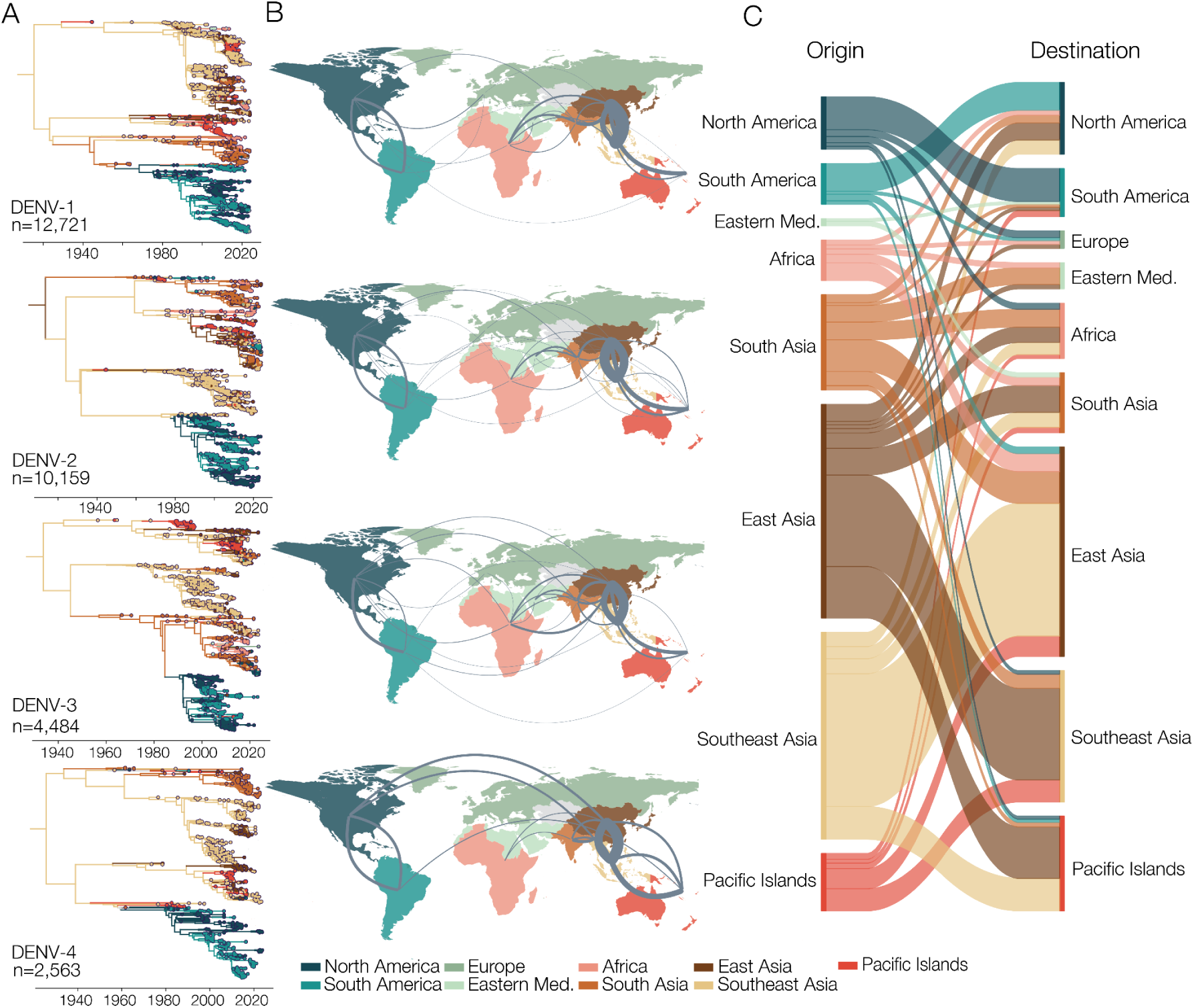
Global phylogeographic analysis of DENV serotypes 1-4. (**A**) Time-resolved phylogeny, colored by inferred location, shows that the global distribution of DENV is highly geographically structured. Estimated root ages DENV-1=1915 (95% HPD=1899-1931), DENV-2=1913 (1880-1934), DENV-3=1934 (1916-1949), DENV-4=1926 (1909-1940); (**B**) Frequency of inferred global virus transitions, width relates to posterior expected number of transitions for each analysis, colored by geographical grouping. Direction is counter-clockwise and thickness of lines relates to frequency of pair over the posterior. Only statistically supported pairs (BF>=3) are shown; (**C**) Relative frequency of inferred virus transition between significant pairs across all serotypes, origin on the left and destination on the right. Same legend used throughout the figure, “eastern med.” refers to the Eastern Mediterranean region.

We found that the inferred root of each serotype was in Southeast Asia (DENV-1, 3 and 4, posterior probability [PP]=64.9%, 67.4% and 50.6% respectively) or East Asia (DENV-2, PP=46.9%), and that the current global lineages of DENV emerged in human populations during the late 1800s to early 1900s (DENV-1 posterior mean=1915 [95% Highest Posterior Density (HPD)=1899-1931], DENV-2=1913 [1880-1934], DENV-3=1934 [1916-1949], DENV-4=1926 [1909-1940]; **Figure 1A**), aligning within the uncertainty intervals of previous estimates (*2*–*5*). We then explored how DENV lineages spread from these emergence points, using Markov jumps between statistically supported pairs of locations (*26*), which we refer to as transitions. We found that most DENV spread is maintained within continents, especially within Asia, accounting for on average 65-75% of the inferred number of between region transitions across the four serotypes (total posterior mean inter-region transitions =191, 238, 176, and 94 for each serotype respectively; **Figure 1B-C**). The two most common pairs of supported virus transitions for each serotype were between East and Southeast Asia: on average, 43% (DENV-1 and DENV-4), 42% (DENV-2), and 36% (DENV-3) of all virus transitions (**Figure 1B-C, S2**).

Our results show that Asia — defined as East, Southeast, and South Asia and the Pacific Islands — is the most important source for global DENV lineage spread: 10 of the 12 most common inter-continental routes across all four serotypes originate in Asia. The most frequent destination is Africa (**Figure 1B-C, S2**). For example, of the 58 possible inter-continental pairs, the most common route for DENV-1 is South Asia to Africa (23% of intercontinental DENV-1 transitions), followed by East/Southeast Asia to Africa.

We infer that Africa is both a source and sink for DENV lineage spread to other regions (**Figure 1B-C, S2**). There are successful introductions (more than one sequence in the dataset, indicating local transmission) from Africa to North America (n=2), South Asia (n=1), and East Asia (n=5). Introductions from South Asia (n=12), East Asia (n=7), and North America (n=2) also have successful circulation in Africa. We observe these despite significant under-sampling of DENV genetic diversity in Africa, indicating its role in the global network may be underestimated (*27*).

Conversely, we show that DENV lineages circulating North and South America are mostly imported from Asia and Africa, but not the reverse (**Figure 1B-C, S2**). Specifically, we estimate that twice as many inter-continental DENV transitions end in the Americas as start there (14.4% vs 7.7%), and the Americas are the destination in well-supported pairs for all Asian regions, Africa, and the Eastern Mediterranean. Finally, both North and South America are well-supported sources of DENV introductions in Europe, aligning with known transmission case studies (*18*, *28*).

### Increasing rate of DENV introductions to the Americas

Our results suggest that — at least with respect to inferred virus spread — the Americas are mostly a sink for DENV lineages from other continents (**Figure 1**). Therefore, we further investigated the source, location, and timing of those introductions, and whether the rate has increased over time (**Figure 2**).

**Figure 2:**
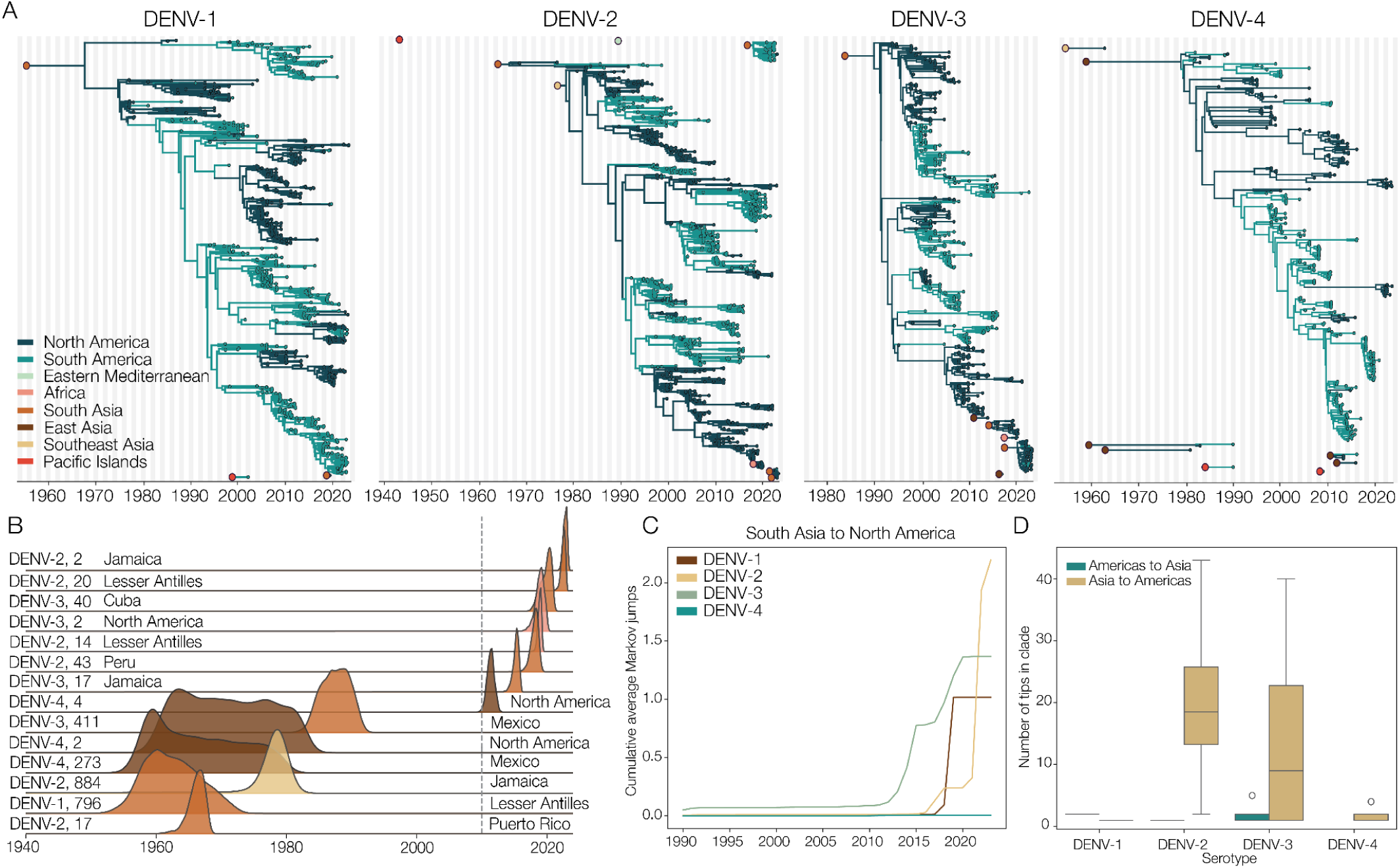
DENV introductions in the Americas. (**A**) Exploded tree view of introductions for each serotype into the Americas from any region from the summary tree, ancestor colored by origin. Each serotype has one dominant clade which still circulates today; (**B**) Probability kernels of non-singleton introduction timing into the Americas across the full posterior distribution. Rate appears to be increasing after ∼2010-2015 following a lull in 2000-2010. Grey dotted line to indicate 2010. Serotype, number of sequences in consensus tree, and inferred origin of location from regional analysis indicated on each line; (**C**) Cumulative mean Markov jumps from South Asia to North America over time by serotype; (**D**) Size of introductions from the consensus tree, with the main American introduction removed for each serotype.

Overall, we estimated 25 introductions into the Americas (unique clades with at least one sequence) in our dataset across the four DENV serotypes (**Figure 2A**). For each serotype, there is a clearly dominant American clade, which arises from introductions into North America (defined geographically as Mexico, Central America, and the Caribbean) during the 1960-90s (DENV-1=1968 [95% HPD: 1962-1974], DENV-2=1979 [1977-1981], DENV-3=1990 [1988-1992], DENV-4=1980 [1978-1981]; **Figure 2B**), similar to previous estimates (*6*, *29*, *30*). Our phylogeographic estimates suggest that the introductions were mostly from Asia (n=22/25), with Africa (n=2) and the Eastern Mediterranean region (n=1) as less common sources. We further estimated that the majority of the introductions were first established in North America (n=18) rather than South America (n=7).

Of the 25 inferred introductions, 14 were successful (included more than one sequence), indicating at least some level of local transmission. We estimated that 12/14 of these introductions originated from Asia (South Asia as the most common individual source, n=8/14) and the vast majority were first established in North America (n=13/14; **Figure 2B**). Further, we inferred that just over half (n=8) of these introductions occurred since 2010, after a quiescent period of ∼20 years since the establishment of the dominant American clades (**Figure 2B**). We also found that the number of transitions has increased since 2010 (**Figure 2C**), suggesting that the actual introduction rate is increasing and not just introduction success. These recent successful introductions were all either DENV-2 or DENV-3, of which half (n=4/8) were from South Asia to North America.

The increasing number of introductions into the Americas since 2010, especially from South Asia to North America, is plausibly due to a bias towards higher sequencing coverage closer to the present. There was a higher proportion of sequences from South Asia in our dataset during 2010-2024 (∼15-20% across the serotypes) relative to 1990-2010 (∼2-6%), while the proportion from North America decreased between 1990-2010 (∼15-23%) and 2010-2024 (∼10-19%; **Figure S3A**). Despite an increase in sequence proportions for all serotypes from South Asia, and a corresponding increase in Markov jumps between all locations from 1990-2010 to 2010-2024 (**Figure S3B**), we only observed an increase in Markov jumps from South Asia to North America for DENV-2 and DENV-3 in recent years (**Figure S3C**). We also tested the association of the sequence proportions from South Asia and North America with the proportions of Markov jumps between these regions using a linear regression (**Figure S3D-G**). We found significant (p<0.001) relationships for DENV-1 (*r*^2^=0.16) and DENV-2 (*r*^2^=0.34), but no relationship for DENV-3 and DENV-4. To explore DENV-2 further, we performed the same phylogeographic analysis using stricter downsampling schemes: 5, 8, and 10 sequences per region per year, which provided a more even number of sequences over time (**Figure S4A**). The two DENV-2 multi-sequence introductions from South Asia that we detected in the original analysis were present in every downsample, although the most recent introduction was present only as a single sequence (but a different sequence in the different downsampling schemes, **Figure S4B-D**). Importantly though, we still detected the increase in transition rate in each of the downsampling schemes (**Figure S4E**). Overall, our rigorous analyses support our finding of an increased rate of DENV-2 and DENV-3 introductions from South Asia to North America in recent years.

Finally, introductions are more likely to transmit locally when spreading from Asia to the Americas compared to the reverse. There are some small introductions from the Americas into Asia that do transmit locally, but these are rare and in general lead to smaller clades than introductions from Asia to the Americas (**Figure 2D**). The same is also true of DENV spread from Africa to Asia (**Figure S5A**), and introductions from Asia are also more successful in Africa than those from the Americas (**Figure S5B**).

### Seasonal synchrony and travel volume drive asymmetrical dynamics of DENV spread

We combined travel, epidemiological, and DENV genetic data to better understand the phylogeographic patterns of DENV spread between Asia and the Americas (**Figure 1, 2**). First, we observed that DENV introductions from Asia to the Americas increased since ∼2010 (**Figure 3A**), which we hypothesized was driven by infected travelers, as the seasonal dynamics have remained relatively consistent over time (**Figure S6A**). The number of travelers from dengue-endemic countries in Asia to the Americas rose considerably from 740,000 in 2000 to a peak of ∼2 million in 2019 (**Figure 3B**). This coincided with a progressively increasing dengue incidence rate in South and Southeast Asia (**Figure S6B**), leading to a higher risk of DENV exposure in travelers from that region. Combining these data, we estimated that the overall DENV travel risk from Asia to the Americas increased over time from 2000 and especially since 2010, interrupted by the global decline in travelers at the start of the COVID-19 pandemic (**Figure 3C**). As incidence measures can be biased within and between regions, we also analyzed data on the number of American travelers diagnosed with dengue during or after travel to Asia from the GeoSentinel network (*31*), which recorded increasing numbers of travelers with symptomatic dengue by site since 2007 (**Figure S6C**). Together, our data suggest that rising numbers of travelers, potentially with more DENV infections, may explain the recent increased rate of introductions in the Americas.

**Figure 3:**
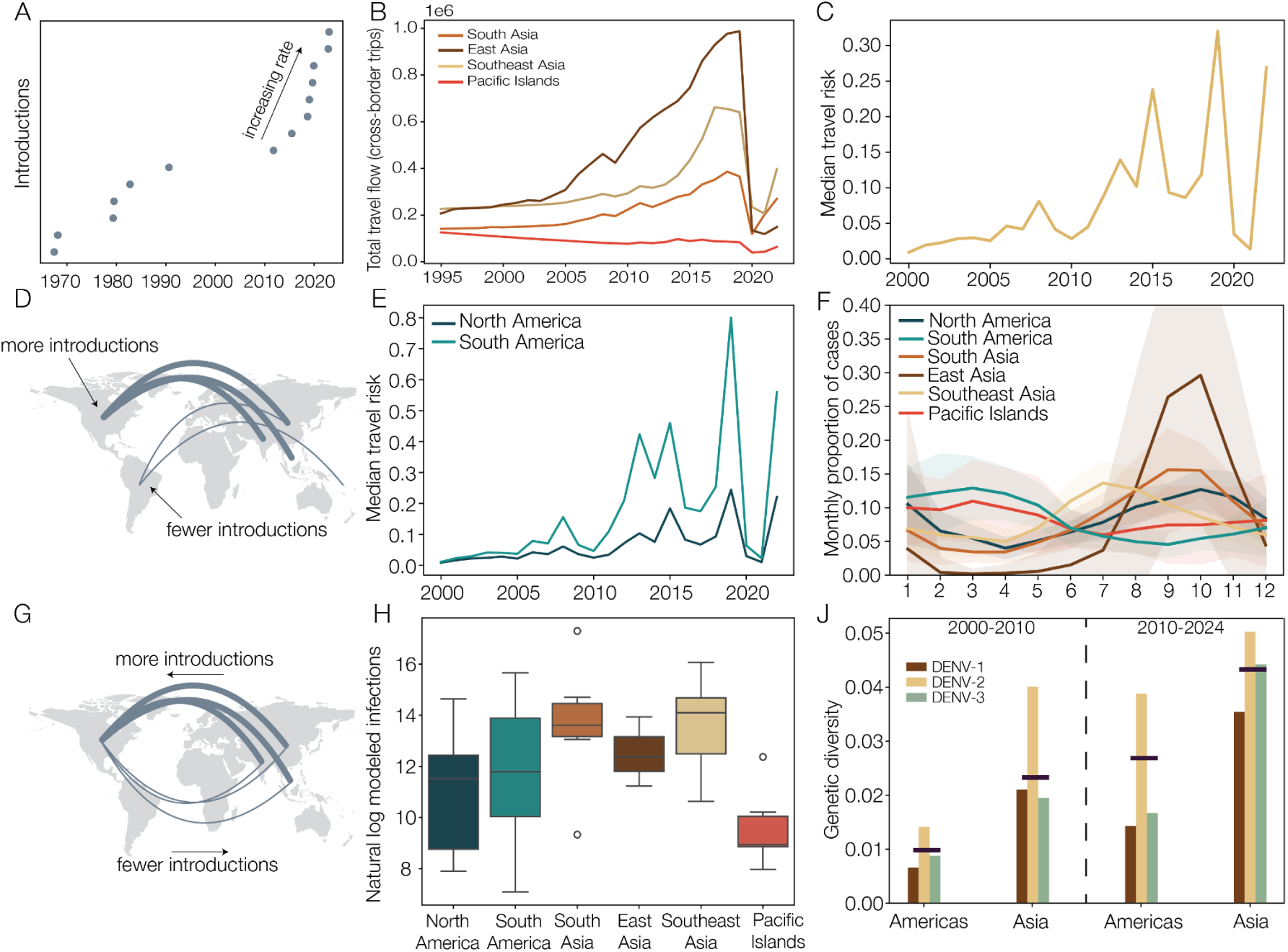
Potential explanations for observed phylogeographic patterns of asymmetric trans-hemispheric DENV spread. (**A**) Schematic showing the timing of introductions into the Americas; (**B**) Total travel flow to both North and South America by Asian region in tens of thousands; (**C**) Median travel risk (air travel passengers ✕ origin dengue incidence rate per person) over time from Asia to the Americas; (**D**) Schematic showing that introductions are mostly successful in North America; (**E**) Median travel risk from Asia to North (dark blue) and South (light blue) America respectively; (**F**) Percentage of cases detected per month in each country, averaged across regions and years, from 2000 onwards. Line is mean and shading is 95% confidence interval; (**G**) Schematic showing that introductions from the Americas are rarely successful in Asia; (**H**) Log modeled infections for each region, from (33). (**J**) Pi diversity of genomic subset for the same regions, split by epoch (2000-2010 and 2010-2024) and serotype. DENV-4 left out due to lack of genomes to generate an even dataset.

Second, we found that most successful DENV introductions from Asia became established in North America (**Figure 3D**) despite estimating a higher opportunity (travel risk driven by a larger volume of travelers) from Asia to South America (**Figure 3E, S6D**). We hypothesized that this is due to differences in onward DENV transmission suitability, and therefore local establishment, at the time of the introductions. We used the proportion of monthly dengue cases reported by each country to show that the dengue seasonality in Asia — especially South Asia — better aligns with countries in North America (South Asia *r*^2^=0.68, Southeast Asia *r*^2^=0.26, East Asia *r*^2^=0.65) compared to South America (South/Southeast/East Asia all negative *r*^2^ values, **Figure 3F**). We also show the same seasonal synchrony between these Northern Hemisphere regions using Index P, a proxy for the basic reproduction number of DENV in mosquitoes based on temperature and humidity (**Figure S6E**). Therefore, at the time when there is a higher DENV infection risk to travelers starting their journey in Asia, there is also a higher risk of onward transmission (success) when infected travelers enter North America relative to South America. Conversely, dengue seasons from South America and the Pacific Islands are synchronous (**Figure 3F, S6E**; *r*^2^=0.36). Despite this being the inferred route of Zika virus spread to the Americas during a period of global emergence (*32*), we hypothesize that the generally small populations and lower incidence rates in the Pacific Islands limits this as a major route for DENV spread into the Americas.

Third, we found that DENV spread between Asia and the Americas is asymmetrical (**Figure 3G**). We hypothesize that the lack of successful DENV introductions from the Americas into Asia is due to a combination of seasonal synchrony, travel infection risk at the origin, and/or virus competition at the destination.

The largest burden of dengue in the Americas is in South America (**Figure 3H**), which has asynchronous DENV seasons with most of Asia (**Figure 3F, and S6E**), prohibiting most transmission between the regions. Spread back to Asia would therefore be expected to start in North America. As short-term travel is mostly symmetrical, the main influence differentiating travel infection risk between any location pairs is the infection risk at the origin. To account for differences in case ascertainment and reporting between regions, including for absolute case numbers (**Figure S6F**) and incidence rates (**Figure S6G**), we used modeled DENV infections derived from force of infection estimates using environmental variables (*33*) and found that infections are on average 13 times higher in Asia than North America (**Figure 3H**). This could indicate that there is a higher traveler infection rate coming from Asia than entering. Further, the longer time period of circulation in Asia results in a higher overall genetic diversity (**Methods**) — and therefore potentially a more complex mosaic of population immunity — compared to the Americas (**Figure 3J**), which has increased in time since the 2000s. While the relationship between genetic diversity and antigenic diversity is complex (*34*), the higher genetic diversity in Asia suggests a more diverse immunological landscape which could make it harder for a lineage from the Americas to invade. DENV spread is also asymmetric between Asia and Africa, likely for some of the same reasons, and collectively helps to explain the unbalanced network of global DENV spread (**Figure 1**).

### Endemic DENV spread within the Americas is associated with travel

We previously detected 14 successful introductions (clades with more than one sequence) into the Americas (**Figure 2**). We estimated the root locations for these introductions using a country-level phylogeography or the sampling location when introductions were only or overwhelmingly sampled in a single location and found that the Caribbean is an important entry point for DENV from outside the Americas. Specifically, we estimate that at least 8 of the 14 introductions originated in the Caribbean, including the Lesser Antilles (n=3; lineages DENV-1V, 2II_A.1.1.1, 2II_F.1.1.5), Jamaica (n=3; DENV-2III, 2II_F.1.1, 3III_B.3.1), Cuba (n=1; 3III_B.3.2), and Puerto Rico (n=1; 2I). Of the remaining introductions, we estimated two originated in Mexico, one in Peru (2II_F.1.1.2), and one had too few sequences to establish a root but was only sampled in Colombia, Puerto Rico, and the Lesser Antilles (**Figure 2B**). The remaining two introductions only contained two sequences each, so we do not attempt to estimate the origin location beyond North America.

To explore the dispersal dynamics in the Americas post-introductions, we conducted a regional phylogeographic analysis on the country level within the region (**Figure 4A**) across the eight sufficiently large introductions that have been sampled since 2000 and in more than one location. An animation of their spread can be viewed here: https://www.youtube.com/watch?v=JAhSp2W15Is. While some introductions into the Americas stretch back into the 20th century, we only discuss results from 2000 onwards in an attempt to lessen the impact of differential sampling effort, which is more pronounced further back in time. We found that while most countries in the Americas are net importers of DENV (i.e., more lineages enter the country than leave, n=14/23 countries with supported transitions) the majority of the countries that are net exporters (sending out more lineages than they receive) are in North America (n=6/9; **Figure 4B**). Nicaragua and Jamaica have the highest proportion of exported DENV lineages across the Americas, although Brazil and Venezuela have the most total transitions originating from them (**Figure 4B**).

**Figure 4:**
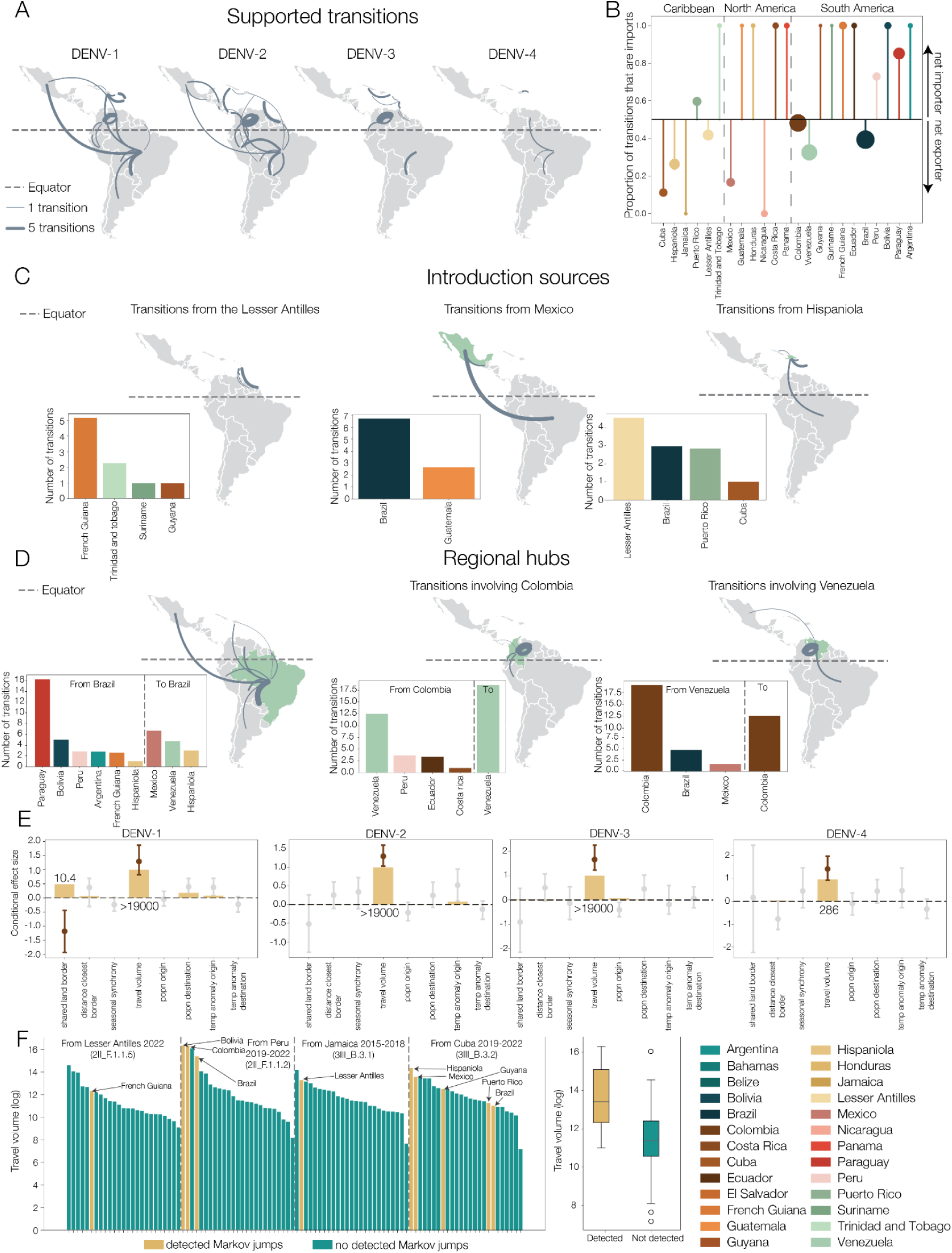
Regional patterns and drivers of DENV spread in the Americas since 2000. For all panels in figure, only pairs where the lower end of the 95% HPD is at least 1 are shown. (**A**) Total transitions by serotype. Direction is counter-clockwise, and line width is by number of transitions, averaged across the posterior across all analyses; (**B**) Proportion of transitions that end in location, colored by country, totalled across all analyses. Horizontal dotted line is at 0.5 (i.e. equal number of transitions starting and ending in a location). Any country above 0.5 is a net importer, and below is a net exporter. Size is proportional to the number of viral transitions. Only pairs where the lower end of the 95% HPD is at least 1 are counted. Vertical dotted lines indicate geographical groupings, indicated at the top of the section; (**C**) Sources of introductions into South America: maps highlighting significant transitions from Lesser Antilles, Mexico, and Hispaniola. Line width is scaled by the total number of transitions across all analyses. Bar charts show the total number of transitions starting in these locations across all analyses and the whole time period; (**D**) Regional hubs: maps highlighting all total transitions across the main introduction for all serotypes since 2000, i.e. established introductions. Line widths and bar charts are the same as above; (**E**) Effect sizes conditional on the predictor being included in the model (dot and whisker) and inclusion probability (bar) for predictors in the phylogeographic model by serotype. Bayes factors are indicated under each statistically supported predictor; (**F**) Travel flow from origin locations of introductions into the Americas for years of lineage persistence (indicated in each box). Blue bars are those without recorded Markov jumps in the analysis, yellow are those with. Box plot indicates all locations together, grouped by whether Markov jumps were detected to each country.

Our phylogeographic analyses further support that DENV lineages spread mostly from North to South America. Out of 21 supported transitions across all introductions (sum of the average from each analysis), 18 originated in North America, 10 of which spread to South America (**Figure S7A-C**). The Caribbean and Mexico play an important role in this process, seeding important regional hubs in South America (**Figure 4C**). Of the three most common transmission pairs, one originated in Mexico (n=7/21, to Brazil) and two originated in the Caribbean (Lesser Antilles [n=5, to French Guiana] and Hispaniola [n=3, to Brazil]). The Lesser Antilles and Mexico were the most common North American origins for spread to South America, responsible for 14/18 total transitions. This mostly one-way flow demonstrates how DENV introductions often originate in North America and then spread to South America, establishing endemic transmission and subsequent outbreaks.

To further understand how DENV lineages are maintained after introduction, we focused on the main American clade for each serotype since 2000 to analyse established DENV circulation. In this analysis, we inferred that DENV spread is mostly contained within South America, making up 9/12 of the top three common between-country lineage transitions for each serotype. We found that Brazil is an important country for lineage spread in South America: it receives approximately half (n=10/21) of transitions from North America when considering all introductions, and is the origin or destination in one of the three most common transitions for the established circulation of DENV-1, DENV-2, and DENV3 since 2000 (**Figure 4D**). Brazil has also seeded significant spread to the highest number of unique locations since 2000 for all serotypes bar DENV-3. Venezuela and Colombia are also important hubs for regional spread, included in half of the most common transitions (**Figure 4D**). Importantly, these three countries are not just sources or sinks but both, indicating centrality within the transmission network, especially Colombia which has approximately equal imports and exports across all introductions (48% of transitions were exports, **Figure 4B**). In general, we found that countries closer to the equator (such as Brazil, Colombia, and Venezuela) were more likely to be both sources and sinks for virus spread (p<0.05, source *r*^2^=0.50, sink *r*^2^=0.35, **Figure S8A**). Finally, the length of time a viral lineage spent in a country was strongly related to country size in square kilometres (p<0.001, *r*^2^=0.85, **Figure S8B**). This result was still evident when Brazil was removed (p<0.001, *r*^2^=0.43).

To estimate the factors defining the network of DENV dispersal in the Americas, we used a phylogeographic model with the transition rates parameterized as a log linear function of potential explanatory covariates (*35*) with each dominant serotype clade. We included all-route travel data (i.e. air, land, and sea) (*36*), distance to closest border, population size, temperature, and seasonality, among other covariates (**Figure 4E**), and found that travel volume was the *only* statistically supported predictor for DENV-2, DENV-3, and DENV-4, and DENV-1 predictors also included a shared land border (**Figure 4E**).

As transmission is associated with travel volume for the dominant DENV clades, we tested whether newer lineage introductions into the Americas mostly spread to countries with higher travel volume, and therefore whether it could be used to predict risk of future spread. In four recent disseminated introductions, areas with higher travel from the inferred origin location tended to have detected DENV transitions (**Figure 4F, S9**). This does not include all high-travel destinations, and we note that there could be undetected spread to other locations that have fewer sequences. Travel volume is, however, clearly an important driver that should be included in future models of risk.

Finally, despite increasing global spread and incidence in the Americas, we found no evidence of increase in average transitions between countries in the Americas over time (**Figure S10**), other than a large increase in 2021 due to the introduction of the lineage DENV-3III_B.3.2. We did, however, find there was an increase in the total number of DENV-2 and DENV-3 transitions, due to new introductions. Therefore each introduction is spreading at a similar rate, but new introductions result in a faster overall spread of DENV throughout the Americas.

## Discussion

Here, we established conclusively that Asia is the source of the past and present global diversity of DENV. Ancestral DENVs emerged from historical sylvatic cycles by the early 1900s before dispersing to Africa, the Pacific Islands, and the Americas throughout the subsequent 100 years (*2*). The timing of these events aligns with major developments in modern transportation, specifically steamships in the late 19th century and affordable air travel starting in the 1970s/80s (*37*, *38*), along with rapid population growth, urbanization, and vector spread (*39*). Asia continues to be the primary source of DENV spread across these travel networks, and lineages are rarely re-introduced into the region. We discovered that DENV lineage spread to the Americas is increasing and mostly follows a specific route: from South Asia to North America before seeding equatorial hubs in South America that maintain regional endemic circulation.

The local establishment of introduced DENV lineages depends on a combination of opportunity (infected travelers arriving) and receptivity (probability of transmission after importation). Opportunity is increasing with the continued growth of global air travel, but success is still rare, with fewer than 20 new DENV lineage establishments in the Americas (magnitudes lower than e.g. respiratory viruses (*40*)) and limited evidence of successful introductions into Asia from the Americas or Africa. We hypothesize this is partially due to seasonal asynchrony between South America and Asia, fewer infected travelers entering than leaving Asia, and to a complex mosaic of local virus genetic diversity and population immunity, making it difficult for introduced lineages to invade. This is particularly pertinent for Asia, where long periods of hyperendemicity and local antigenic evolution lead to higher competition (*34*, *41*). Even when all else appears equal, introductions have had different success. In the mid-1900s we detected sputtering introductions around the same time that the dominant clades for each serotype were established; and in recent years, one DENV-3 introduction in the Caribbean remained mostly confined to the Caribbean (3III_B.3.1) (*18*) while a second, 3III_B.3.2, rapidly emerged across the Americas and Africa (*18*, *42*–*46*). There are therefore complex factors around success (*47*), including population immunity, environmental suitability, and human population mixing that must be further investigated to be able to predict the likelihood of future DENV lineage establishment, all while the opportunity continues to increase.

Despite a higher number of travelers entering South America from endemic regions, we discovered that North America, especially the Caribbean Islands and Mexico, are the primary entry point for DENV introductions in the Americas. We hypothesize this is likely because Mexico and the Caribbean are in the Northern Hemisphere, therefore aligning better with dengue seasonality in Asia (*48*). There are several other reasons why invasion success could be higher here, such as the human and mosquito ecology (dense, coastal cities where *Aedes aegypti* thrive), immunological landscape (islands can experience sharp epidemic waves followed by quiescent periods that can generate large cohorts of susceptibles), and climate (sufficient temperature for mosquitoes across most of the year) (*49*–*51*). Specifically for the Caribbean, post-colonial cultural connections and overseas territories result in many Caribbean islands having disproportionately high travel volumes to Asia, Pacific, and Indian Ocean islands, as well as to South America. One of the DENV lineages that we described here (2II_F.1.1.5) spread from South Asia to the French Lesser Antilles, and then to French Guiana in South America, Réunion in the Indian Ocean, and mainland France in Europe (*28*). Therefore, enhanced DENV genomic surveillance in the Caribbean, both locally and via travelers (*18*), would help with the earlier detection of emerging lineages in the Americas.

From North America, we showed that onward spread to equatorial countries in South America, namely Brazil, Colombia, and Venezuela, fuels endemic DENV circulation throughout the Americas (*6*, *52*). This network of regional DENV spread is primarily driven by human travel (*30*, *53*). We further showed that onward spread of a new DENV lineage from the introduction source in the Americas is mostly predictable by travel volumes, which could have major applications for constructing a formal method to forecast the dynamics of introduced lineages. In recent years, many large dengue outbreaks in the Americas have been associated with new lineages introduced from Asia. For example, the introduction and emergence of lineage 2II_F.1.1.2 (also referred to as the "cosmopolitan" genotype) was associated with the explosive 2024 outbreaks, especially in Brazil (*54*–*58*). In parallel, lineage 3III_B.3.2 was introduced into Cuba and rapidly became the dominant lineage across the Caribbean and parts of Central and South America, driving the sudden resurgence of DENV-3 after a decade of quiescence (*18*, *42*–*45*). Tracking the increase in DENV introductions is essential because emerging lineages can rapidly spread through the Americas’ travel network and drive major outbreaks.

Our analyses are dependent on the currently available DENV genomic data. Specifically, we are unlikely to capture all successful introductions in all regions and timepoints, particularly in countries with lower sequencing capacity and pre-2000 when there was much lower sequencing coverage. We downsampled the existing dataset to reduce the impact of globally heterogeneous sampling, but this means we may not detect every introduction that is present in the full genomic dataset. We also note that our travel risk estimates are dependent on DENV infections being captured by different surveillance systems, and that GeoSentinel infected travelers data includes changes in diagnostic approaches over the study period.

To fully understand changes in global DENV transmission, there are three major missing components. The first is the ability to predict antigenic diversity from genomes (e.g. (*59*), thereby connecting genomic distance to antigenic distance. This would help us to understand a component of differential success of introductions in the Americas, and why American/African introductions into Asia are mostly unsuccessful. Second, Africa clearly plays an important role in the global network (*27*), but it is undersampled and so we almost certainly underestimate its impact. It should therefore be a priority to set up additional dengue surveillance across the continent. Finally, we must understand the epidemiology of DENV in South Asia, especially India. South Asia is a common source for DENV in the Americas, but data are relatively sparse. We expect this under-reporting explains the discrepancy between most introductions originating in South Asia and a non-exceptional travel risk (*60*). Additional data from South Asia would help to better quantify the proportion of incoming DENV lineages that are successful in establishing transmission in the Americas, and make global transmission estimates more accurate — e.g., for influenza viruses, the role of India as a source population was under-estimated until genomic data was routinely collected (*61*). Ultimately, we must be able to understand what new introductions to different regions mean epidemiologically for disease risk, and how this will vary with varying population immunity and climate change.

Overall, we characterized the complex, asymmetric, and dynamic network of global DENV spread, highlighting the importance of a global DENV genomic surveillance system. Our results indicate increasing inter-regional spread, leading to a higher amount of cross-country virus dispersal in receiving regions. While this is partly due to apparent increases in dengue incidence in source populations, which is exacerbated by climate change (*62*), longer-term patterns of spread are mostly governed by travel patterns, climate synchronization, and (perhaps) local competition. Dengue is a disease of industrialized human travel, and our results help to understand how our increasingly connected world can lead to devastating local outbreaks.

## Methods

### Full genomic dataset

All genomes with either over 70% coverage (n=16,348) or which were complete envelope regions (1479-1485 base pairs in length, depending on serotype, n=16,093) were downloaded from NCBI Genbank on 2024-04-11. Both whole genome sequences (WGS) and envelopes (E) were used in all analyses presented in this paper, other than where stated. The most recent dates for each serotype are the end of 2023 or the start of 2024.

We included partial genomes as their inclusion resulted in increased data for some countries (e.g. China, Tanzania), and the inclusion of some countries which would otherwise have been excluded (e.g. Madagascar for DENV-1). To test that the inclusion of E sequences would not impact temporal estimates, we took a subsample of 1000 random WGS throughout the time period, and artificially downsampled half of them to their E sequences only to match the proportion of E sequences in the full dataset. We then conducted a joint phylogeographic analysis on the WGS and the E sequence dataset. We found that the root of the tree, the mean and standard deviation of the rate of evolution, and location clock rate were compatible between both analyses (i.e. had overlapping 95% highest posterior density [HPD] intervals). We therefore concluded that, similarly to Ebola (*63*) and Rabies (*64*) viruses, geographical signal was improved by adding partial genomes; but that over the larger time period and with sufficient WGS to provide additional information, temporal signal was not compromised.

Sequences were serotyped based on the similarity to reference sequences for each serotype, and sylvatic or “Unassigned” sequences removed (genotypes 2VI and 4VI for DENV-2 and DENV-4 respectively, accession numbers KR919820, EF457905, and FN825674 for DENV-1, and KT452800, MW945427, and OM258630 for DENV-3). We also removed sequences without any location information, and identical sequences from the same day and country (n=607) to reduce dataset size and increase inference efficiency.

We returned to publications to supplement date and location information where possible. Our aim was to include genomes from as many locations as possible but still retain computational efficiency in estimating phylogenies, so we aimed to reduce the number of sequences which would require additional inference on their dates. For locations with fewer than 100 sequences in total, we kept all sequences regardless of date information. For locations with more than 100 sequences, if fewer than 10% were missing dates, these sequences were removed. (n=475 across all serotypes). Sequences which had locations known to not be endemic for dengue (e.g. France, USA, Ireland) were examined, and assigned to traveler origin or removed. French overseas departments (e.g. Réunion, Martinique etc.), Puerto Rico, and Hawaii were treated as separate locations to France and the USA for ease of interpreting geographical and climatic patterns. Finally, sequences with uncertain locations (e.g. KP406801, which is listed as “India/Singapore”) were removed.

We used this dataset as input for molecular clock outlier investigation in TempEst (*65*). Sequences which were passaged, very difficult to align, and molecular clock outliers were removed.

The final dataset sizes for each serotype are 12,721 (DENV-1), 10,159 (DENV-2), 4,484 (DENV-3) and 2,563 (DENV-4).

### Rate analysis

We conducted a (log) marginal likelihood estimation analysis to check whether a strict clock model or one allowing rate variation was a better fit for our data.

We subsampled the dataset to up to 1000 whole genomes per serotype by taking up to a maximum number of sequences per year. We preferentially used those with complete dates, and topped up samples with incomplete dates where necessary, and used all sequences in a year if there were not enough sequences. We then used the GTR nucleotide substitution model (*66*) and the skygrid coalescent model with the Hamiltonian Monte Carlo (HMC) transition kernel and gridpoints every five years (*67*, *68*). We ran two independent chains to check for convergence in Tracer (*69*).

We compared stepping-stone sampling results (*70*), and found that the relaxed clock was a better fit for our data than the strict clock model (**Table S1**).

### Global downsample and discrete trait analysis

We started with global areas based on those defined by the World Health Organization (WHO), and we did not change Europe and the Eastern Mediterranean regions. We split the Americas into North and South America, and changed the definitions for Asian regions to separate South Asia, Southeast Asia, East Asia and the Pacific Islands. See **Table S2** for a list of countries in each region.

We downsampled based on these regions to increase computational efficiency and to decrease sampling bias between regions where possible. We selected a number of genomes per region per year so that South America and Southeast Asia were similarly represented in the downsampled dataset as two regions of major DENV circulation. The maximum number of sequences per region per year that was used was 20 (DENV-1 and DENV-2), 15 (DENV-3), and 10 (DENV-4). We did this instead of using case data to downsample because comparing DENV surveillance efforts over the whole globe and a century of transmission introduced additional biases that we could not control for. We therefore concluded that attempting to harmonise based on existing knowledge of DENV transmission reduced this bias.

To select these sequences, we first prioritised whole genomes with complete dates, then with months, then only years. The same procedure was then repeated with envelope regions until the maximum number per year was reached. The final dataset sizes for the global geographic analysis were 2492 (DENV-1), 2625 (DENV-2), 1570 (DENV-3) and 823 (DENV-4). See **Figure S1** for the distribution of sequences over time.

In the interests of efficiency, we split the analysis into the tree topology estimation and a subsequent discrete phylogeographic analysis using a forward-in-time (FIT) asymmetric continuous-time Markov chain (CTMC) model. We used an uncorrelated relaxed clock with an underlying lognormal distribution and HMC transition kernel (*71*), based on results obtained from the analysis above, using tipdate sampling (*72*) on sequences with only year information. We used a skygrid tree prior (*67*) with an HMC transition kernels (*68*) with one gridpoint per year, a GTR nucleotide substitution model (*66*), and a gamma distributed rate heterogeneity model with four rate categories (*73*). We ran two to three chains for each serotype, removing 10-25% for burn-in.

We then took 1000 trees from across the post-burn-in posterior from the above analysis to run the FIT phylogeographic analysis (*74*) in BEAST X (*75*, *76*), across 9 locations (“Southeast Asia”, “East Asia”, “South Asia”, “Pacific Islands”, “North and Central America and the Caribbean”, “South America”, “Africa”, “Europe”, “Eastern Mediterranean”), using Bayesian stochastic search variable selection (BSSVS). Two chains each of 5 million states were run, 10% removed for burn-in. Introductions into the Americas were defined at the American node closest to the root that had a parent outside of the Americas. There were no nested introductions (i.e. lineages which leave and re-enter the Americas) detected.

For all FIT phylogeographic analyses “transitions” are inferred Markov jumps (*26*) between discrete states which have been averaged across the posterior.

All time-resolved phylogenies shown in this paper are highest independent posterior subtree reconstruction (HIPSTR) trees which summarise the posterior tree distribution (*77*).

### Downsample for regional FIT analysis on main American clade

For the analysis on the main clade for each serotype in the Americas, we needed to generate a new downsampled dataset to minimise sampling bias on a regional scale. We started by adding sequences, which we had previously removed to reduce the number of American sequences on a global scale, back to the main clade for each serotype. We did this by finding the most recent common ancestor (MRCA) of the main clade in the maximum likelihood tree of the complete (i.e. un-downsampled) dataset and took every sequence that was downstream of that node that was sampled in the Americas. We validated this by checking that the timing of the MRCA in the downsampled and non-downsampled datasets had overlapping HPDs. This therefore gave us a complete dataset for the sections of interest in each tree.

Next, we downsampled using estimated DENV infections based on a modeled force of infection (FOI) (*33*). FOI is the per capita rate at which a susceptible individual becomes infected with a given pathogen. The authors in (*33*) used this to estimate global DENV burden at fine spatial resolutions, in an attempt to overcome biases in reporting within and between countries. We therefore used this modeled estimated number of infections to downsample rather than reported cases.

Not every country has an appropriate modeled FOI underlying its infection number. This includes the contiguous USA, where the value is based on a single datapoint from Texas, and most of our sequences were from Florida. Further, as the USA is a large country that has only sporadic DENV transmission in a small number of areas, non-genomic data such as number of travelers and climate indicators would primarily be from parts of the country with no dengue, and skew results. We therefore removed sequences from the continental USA. The other missing locations tended to be Caribbean islands. To solve this, and to reduce the number of states with low numbers of sequences in the DTA, we grouped the Lesser Antilles together (other than Trinidad and Tobago which is only sometimes included in this group) and averaged the available FOIs.

We calculated the percentage of the total modeled infections in the Americas that each country represented. This gave us the relative contribution of infection to the region for each country, and therefore the expected relative contribution of genomes to the total genomic dataset. We multiplied the total number of sequences per location per year by this percentage, and rounded to the nearest whole number to get the expected number of genomes. If the expected number of genomes was rounded down to zero, a single sequence was included. These sequences were picked randomly from across the year. This resulted in a dataset where only Mexico (DENV-2, DENV-3, and DENV-4), and Brazil (DENV-3) were more than 5 percentage points different from the desired percentage. We also tried a year agnostic approach, but this resulted in a larger difference between the actual percentage of sequences in the dataset and the percentage of regional FOI for the same countries. The final dataset size for the main introduction American downsample was 936 (DENV-1), 756 (DENV-2), 284 (DENV-3) and 265 (DENV-4).

### Other American clades

To estimate the dynamics of other introductions into the Americas, we performed the same procedure as described above to add in previously downsampled sequences to all introductions. We then took introductions of three or more sequences from the full dataset for inclusion in the analysis (DENV-1 n=0, DENV-2 n=5, DENV-3 n=2, DENV-4 n=1). For introductions of more than 50 sequences, we downsampled by taking 10 sequences per location per year, keeping all locations to accurately estimate the root location. Clades of fewer than 50 sequences were not downsampled.

### Non-genomic data

We obtained passenger flow data from the Global Transnational Mobility Dataset (*78*). This combines air traffic, tourism, land and sea traffic, and migration statistics to obtain an estimate of all travel across borders. Our aim was to investigate travel flow between regions with local DENV transmission. As defining DENV endemicity is challenging, we used a series of criteria to define these regions. First, we only included countries with genomic sequence data. Second, we removed the USA, Australia, South Korea, Japan, and Chile from this dataset as DENV transmission in these countries is sporadic and due to virus introductions from elsewhere, while their high volumes of international air travel could skew our results. We retained China and Argentina even though DENV is not endemic throughout either country, because most of their populations reside in areas with consistent DENV transmission (*79*, *80*).

Population data was extracted from Worldometer, and missing years were interpolated using the population growth percentage. Land area in kilometres squared was also obtained from Worldometer. Dengue case data was extracted from OpenDengue (*81*). Seasonal synchrony was estimated using the percentage of case counts that occurred in each month for each country (*60*, *81*), as this should be internally consistent with how countries measure and report DENV. We also confirmed this with index P, an estimate of viral reproductive fitness in mosquitos, obtained from (*82*). We used an estimate of temperature anomaly as a predictor in the regional-scale phylogeographic model with covariates, as described previously (*49*).

For each parameter that had to be averaged into epochs and groups, we first averaged across the groups, so there was one value per region per year, and then across years in the epoch.

Data on returning travelers with dengue were obtained from GeoSentinel, a global clinician-based sentinel surveillance network of specialised travel and tropical medicine sites across six continents (https://geosentinel.org/sites).

### Regional discrete trait analysis

We ran FIT phylogeographic models on the largest introduction for each serotype, which is one genotype (DENV-1V and DENV-2III) or major lineage (DENV-3III_C and DENV-4II_B (*83*)). The analyses were once again split into tree topology estimation and the FIT analysis. The same parameters as the global dataset were used, and two MCMC chains were run for each serotype of 50-110 million states with 10-20% removed for burn-in. Similarly, 1000 posterior trees were used as input for each FIT analysis.

To include time-varying predictors and identify potential changes in regional dynamics, we used a time-inhomogeneous model for the rate and predictor matrices (*84*). In total, there were six epochs, one every five years from ∼2024 to 2000, and then one epoch for all years before 1999. The oldest epoch parameterises viral transitions using a standard FIT CTMC without predictor matrices. The most recent epoch has slight variation in length for different analyses due to slight differences in the time of the most recent tip, but having the same change-points between analyses was prioritized (over having the most recent epoch be exactly five years) in the interest of comparability.

For each epoch other than the oldest, we included an approach to parameterize transition rates between locations (*35*). Some predictors included in this were static over time (shared land border, distance to the closest border); and others were dynamic (seasonal synchrony, travel volume, population and temperature anomaly). For dynamic predictors, values were averaged across the available years in the epoch. All predictors were log-transformed and standardised. After preliminary analysis showed that inclusion probabilities did not vary much between epochs, coefficients and indicator values were shared across all epochs with predictors to maximise signal.

Some predictors did not have values for every location (e.g. temperature anomaly). In cases where more than 50% of locations in a group were missing a predictor value, this was treated as missing for the group. For missing predictors, we leveraged a new BEAST method to sample missing covariate values from their full conditional posterior distribution (*85*). If they were missing for less than 50% of locations, an average was taken to represent the group.

For smaller introductions, we ran a country-level DTA to estimate the introduction location and subsequent geographical dynamics. These introductions were sub-lineages of 2II_A.1.1.1 (n=1), 2II_F.1.1 (n=3), 2I, 3III_B.3 (n=2), and 4II_A.1.2 (n=1) (*83*). Some introductions were too small to be run independently (e.g. the DENV-4 introduction), and others were only documented in one location (e.g. a DENV-2 introduction that stayed in the Lesser Antilles). We therefore ran five independent analyses on three DENV-2 introductions and two DENV-3 introductions. We used an uncorrelated clock with HMC inference and tip date sampling as in the global analysis, and the skygrid population model with an HMC transition kernel (*67*). We ran two independent chains for each analysis for 100 to 500 million states, and removed 10-20% of states for burn-in.

Introductions into South America from North America were defined as South American nodes with North American parents that had more than two sequences, and a posterior probability of origin location of at least 50%. Lineages which left South America and returned later are counted as distinct introductions.

To calculate the percentage of transitions that end in a country across all American introductions, we first averaged the number of transitions that start or end in a specific country across the full posterior of each location. Then, we added these all together between introductions, and calculated the proportion based on these totals between all locations.

### Genetic diversity

To explore differences in genetic diversity between the Americas and Asia, we took 100 whole genome sequences across each epoch (2000-2010 and 2010-2023), with at least five sequences in every year. We did not include DENV-4 because there were not at least five whole genome sequences in each year from 2000-2023 from both Asia and the Americas.

We used snp-sites (*86*) to generate an invariant Variant Calling Format (VCF) file from the multi-sequence alignment of each serotype-epoch pair, which we zipped and indexed with samtools (*87*). We then calculated genetic diversity using **π** diversity (*88*), the average per site polymorphism, using pixy (*89*) which takes missing data into account. Only positions with exclusively A, T, C or G were used in the calculation.

## Acknowledgements

We thank N. Nosenko for her thoughts on GIS mapping, C. Brookson, S. Good, and Y. T. Chew for their guidance on risk modeling, P. Carlton, P. Jack, S. Taylor, and V. Parsons for technical support, GeoSentinel network for providing the travel data used in this analysis, and all those who have generated and shared genome sequence data which makes work like this possible. This publication was made possible by the National Institute of Allergy and Infectious Diseases of the National Institutes of Health (NIH) under Award Number DP2AI176740 (NDG). VH and GB acknowledge support from the Research Foundation - Flanders ("Fonds voor Wetenschappelijk Onderzoek - Vlaanderen," G098321N). VH acknowledges support from the Wellcome Trust (327522/Z/25/Z). VH, SD and GB acknowledge support from the European Union Horizon 2023 RIA project LEAPS (grant agreement no. 101094685). GD has received funding from the Research Council of Lithuania, Lithuania under the EMBO Installation Grant programme under grant No 5305. HR has received support from the National Science Foundation (DGE-213984). NRF acknowledges support from the Wellcome Trust Dengue and Zika Immunology and Genomics Multi-Country Network (DeZi Network; 316633/Z/24/Z). M.U.G.K. acknowledges funding from The Rockefeller Foundation (PC-2022-POP-005), Health AI Programme from Google.org, the Oxford Martin School Programmes in Pandemic Genomics & Digital Pandemic Preparedness, European Union’s Horizon Europe programme projects MOOD (#874850) and E4Warning (#101086640), Wellcome Trust grants 303666/Z/23/Z, 226052/Z/22/Z & 228186/Z/23/Z, the United Kingdom Research and Innovation (#APP8583), the Medical Research Foundation (MRF-RG-ICCH-2022-100069), UK International Development (301542-403), the Bill & Melinda Gates Foundation grants (INV-063472, INV-090281, INV-103122) and Novo Nordisk Foundation (NNF24OC0094346). OJB is supported by a United Kingdom Medical Research Council grant (MR/V031112/1) and acknowledges funding from the AXA Research Foundation (title: “Pioneering a global observatory for dengue outbreak early warning”) which also supports HJ. SD acknowledges funding from the *Fonds National de la Recherche Scientifique* (F.R.S.-FNRS, Belgium) and from the University of Brussels (ULB, Belgium) internal fund. PL acknowledges support by the Research Foundation - Flanders (‘Fonds voor Wetenschappelijk Onderzoek - Vlaanderen’, G010326N and G051322N). PL and GB acknowledge support from the Internal Funds KU Leuven (Grant No. C16/26/012). GB acknowledges support from the DURABLE EU4Health project 02/2023-01/2027, which is co-funded by the European Union (call EU4H-2021-PJ4) under Grant Agreement No. 101102733. GeoSentinel is co-funded through a Cooperative Agreement between the US Centers for Disease Control and Prevention (CDC) and the International Society of Travel Medicine (1 U01CK000632-01-00). The findings, contents, and conclusions in this report are those of the authors and do not necessarily represent the official position of the NIH, CDC, European Commission, or other funders.

## Notes

### Competing Interest Statement

The authors have declared no competing interest.

