## Supplementary figures for "Phylodynamic analysis reveals asymmetric and accelerating global spread of dengue virus"

| Serotype | Strict clock | Relaxed clock |
|---|---|---|
| DENV-1 | -177812.20 | **-177223.80** |
| DENV-2 | -185097.73 | **-184467.38** |
| DENV-3 | -156500.89 | **-155086.08** |
| DENV-4 | -152683.31 | **-151443.03** |

***Table S1: Results of stepping-stone sampling analysis*** *to assess which clock model to use based on fit to the data. For all serotypes, an uncorrelated relaxed molecular clock with an underlying lognormal distribution fits the data better than a strict clock.*

***Table S2 Category that each country with DENV sequence data is assigned to***

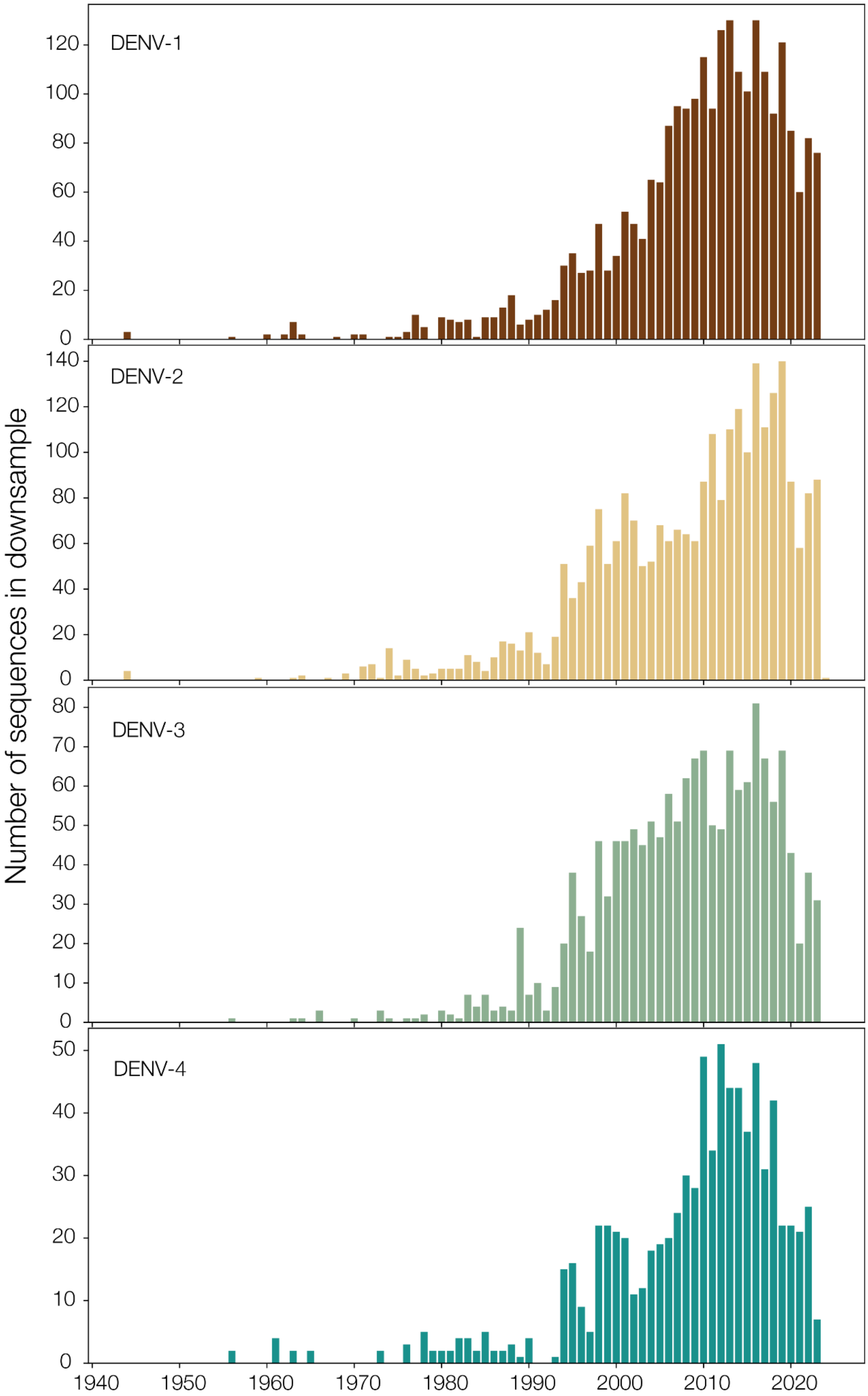


***Figure S1: Dataset used for global phylogeographic analysis.*** *Shows number of sequences in the downsample per year per serotype.*

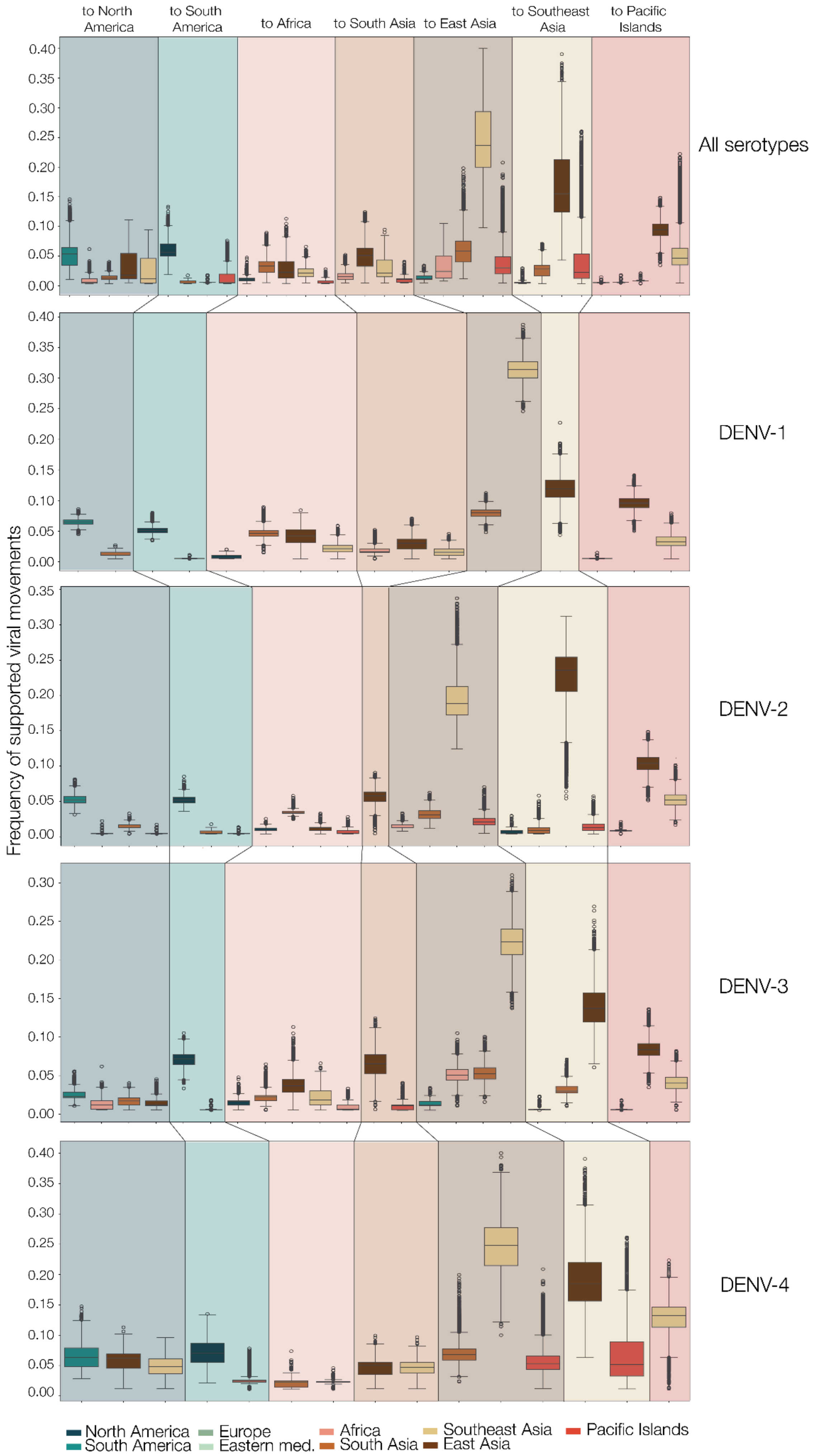

to North America
to South America
to Africa
to South Asia
to East Asia
to Southeast Asia
to Pacific Islands
All serotypes
DENV-1
DENV-2
DENV-3
DENV-4
Frequency of supported viral movements
0.00
0.05
0.10
0.15
0.20
0.25
0.30
0.35
0.40
North America
South America
Europe
Eastern med.
Africa
South Asia
Southeast Asia
East Asia
Pacific Islands

***Figure S2: Virus transitions in global analysis****. Frequency of significant virus transitions for all serotypes together (top row) and by serotype. Boxes are colored by origin location and grouped by destination location, indicated at the top of each section and by background color. Same legend used throughout the figure.*

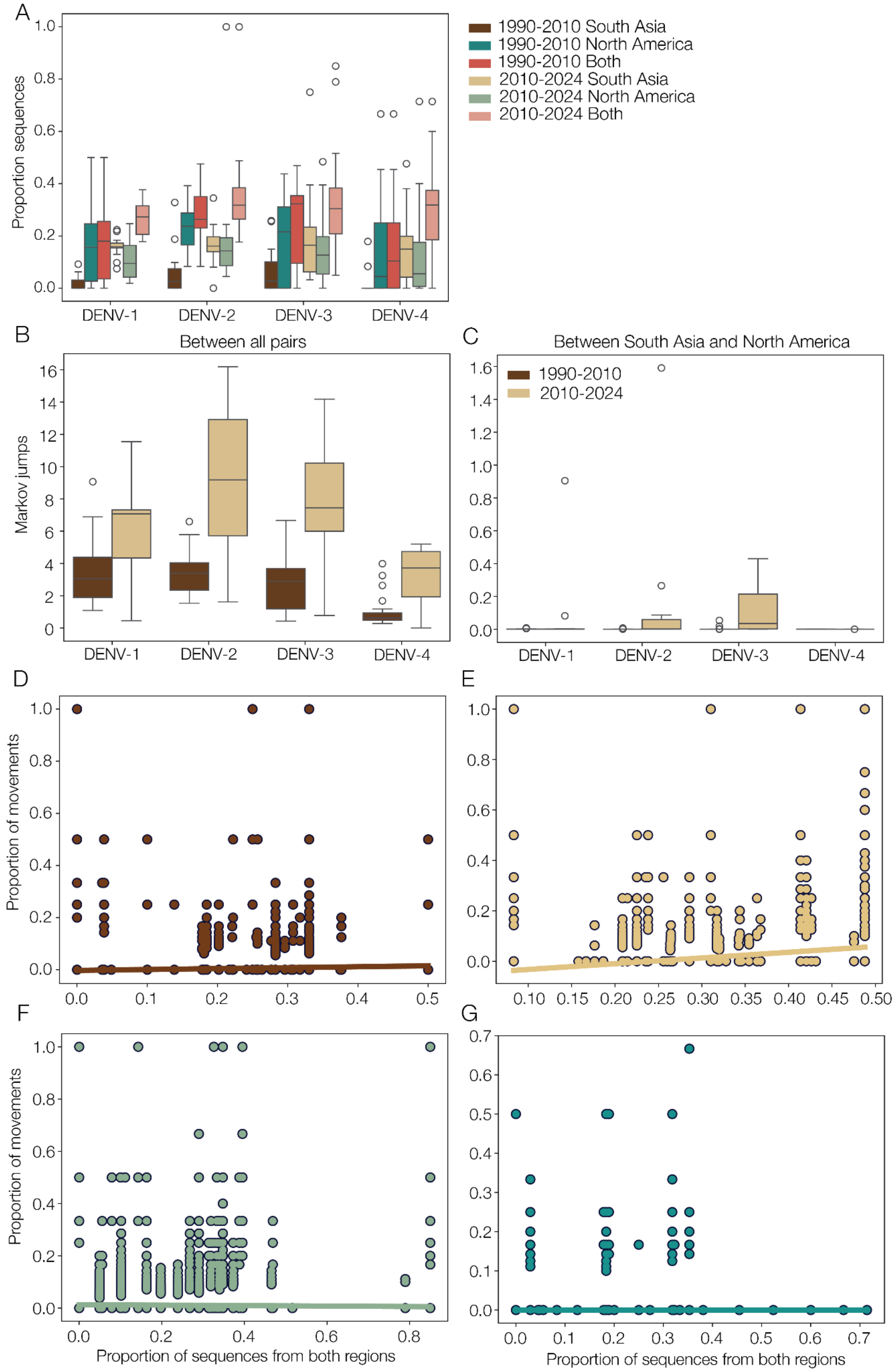


***Figure S3: Validation of increase in introductions**. (**A**) Proportion of sequences in time periods 1990-2010 and 2010-2024 of North America, South Asia and both combined; (**B**)*

*Absolute Markov jumps in each period between South Asia and North America for each serotype and (**C**) all jumps. Absolute jumps between all locations increase for all four serotypes, but on DENV-2 and DENV-3 for South Asia to North America; **D**-**G**) Linear regressions of proportion of Markov jumps that are South Asia to North America against proportion of sequences from both regions for serotypes DENV-1-4 respectively. R-squared DENV-1 = 0.159 (p<0.001), DENV-2 = 0.34 (p<0.001), DENV-3 = -0.021 (p<0.001), DENV-4 0.0014, (p=0.4).*

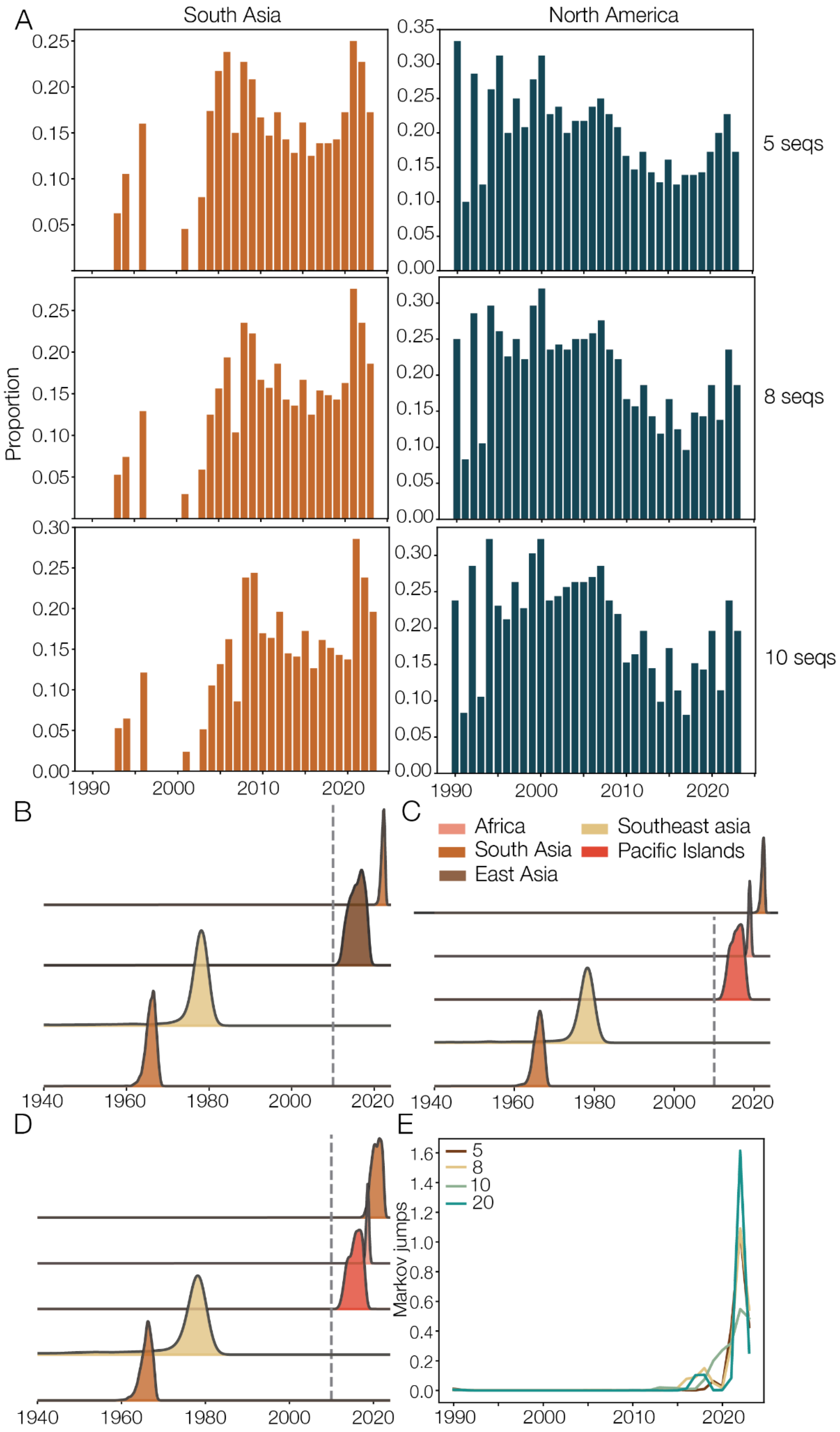

A
South Asia
North America
5 seqs
8 seqs
10 seqs
Proportion
B
C
Africa
South Asia
East Asia
Southeast asia
Pacific Islands
D
E
Markov jumps
5
8
10
20

***Figure S4: Additional DENV-2 downsamples to explore apparent increase in introductions.*** *(**A**) Proportion of sequences in South Asia and North America for downsamples with 5 sequences, 8 sequences, and 10 sequences per year. Kernels of introduction times into the Americas for 5 sequences (**B**), 8 sequences (**C**) and 10 sequences (**D**). Colored by inferred origin. (**E**) Markov jumps over time from South Asia to North America for each downsample compared to original analysis (20 sequences per year).*

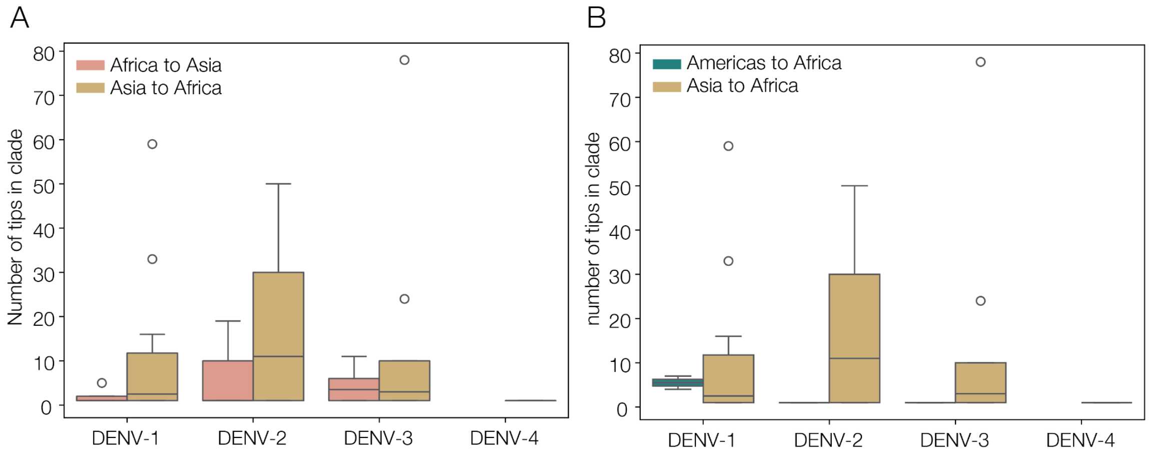


***Figure S5: Differential introduction opportunity and success.*** *(**A**) Tip number on consensus tree of introductions from Africa to Asia (pink) and vice versa (yellow). (**C**) Tip number on consensus tree of introductions into Africa from the Americas (blue) and Asia (yellow).*

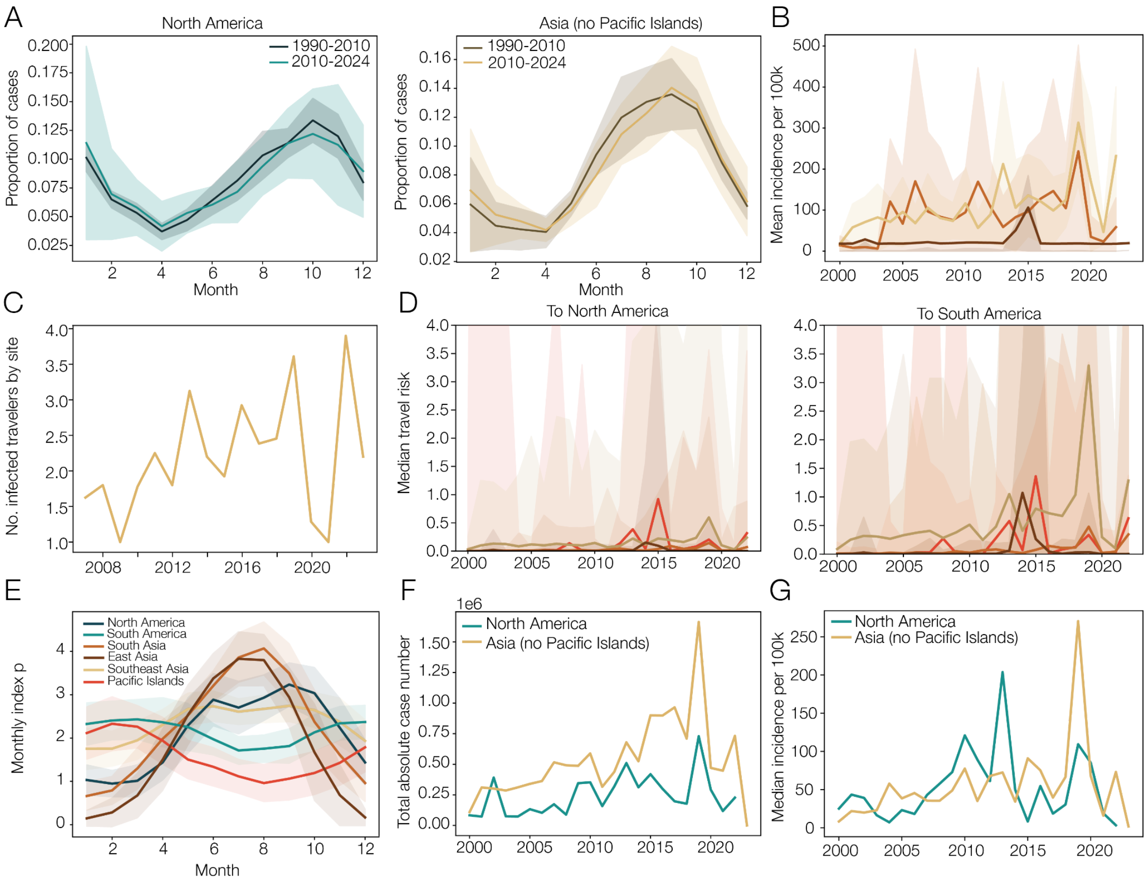


***Figure S6: Contributors to global dynamics.*** *(**A**) Seasonality by proportion of cases per month, averaged across years by epoch. North America shown on the left, East/South/Southeast Asia averaged together on the right; (**B**) Mean dengue incidence rate (cases per 100,000 population by region in Asia, shaded areas indicate 95% CI; (**C**) Number of infected travelers from the Americas diagnosed during or after travel to Asia, normalised by the number of reporting sites per year; (**D**) Median travel risk by Asian region to North America (left) and South America (right). Shaded areas indicate 95% confidence interval (**E**) Seasonality for each region as shown by index P, a measure of viral reproductive fitness in the mosquito host population. (**F**) Total absolute cases in millions for synchronised American and Asian regions. (**G**) Incidence per 100,000 population for North America (blue) and Southeast/east/south Asia (yellow) - i.e. the synchronised regions; Legend for regions the same, other than where indicated, can be found in panel E.*

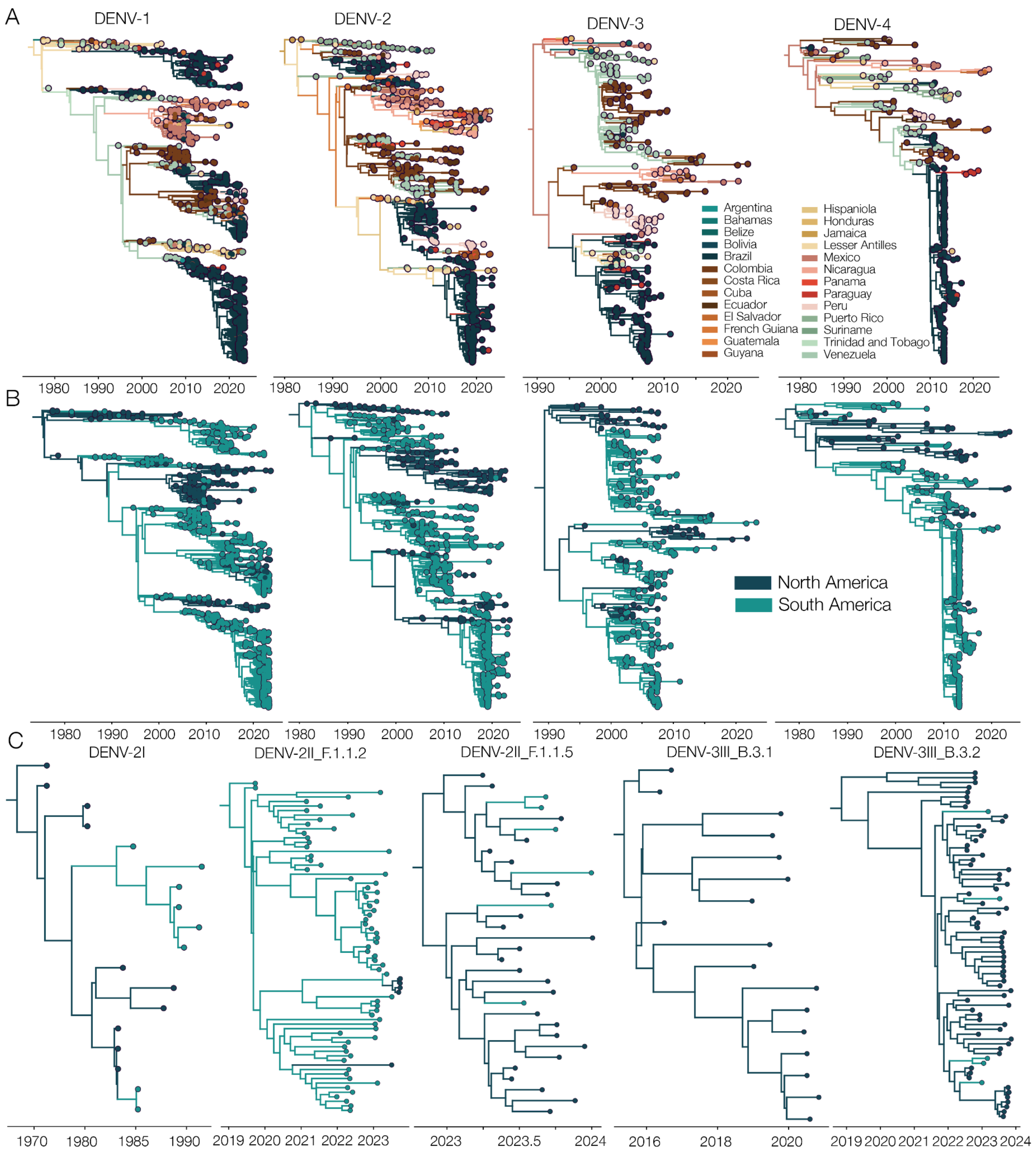


***Figure S7: Regional spread of DENV**. (**A**) Time-scaled phylogenies of each of the main introductions per serotype, colored by country; (**B**) The same phylogenies colored by North America vs South America; (**C**) Time-scaled phylogenies of minor introductions, labelled by the lineage they are part of, and colored by North vs South America.*

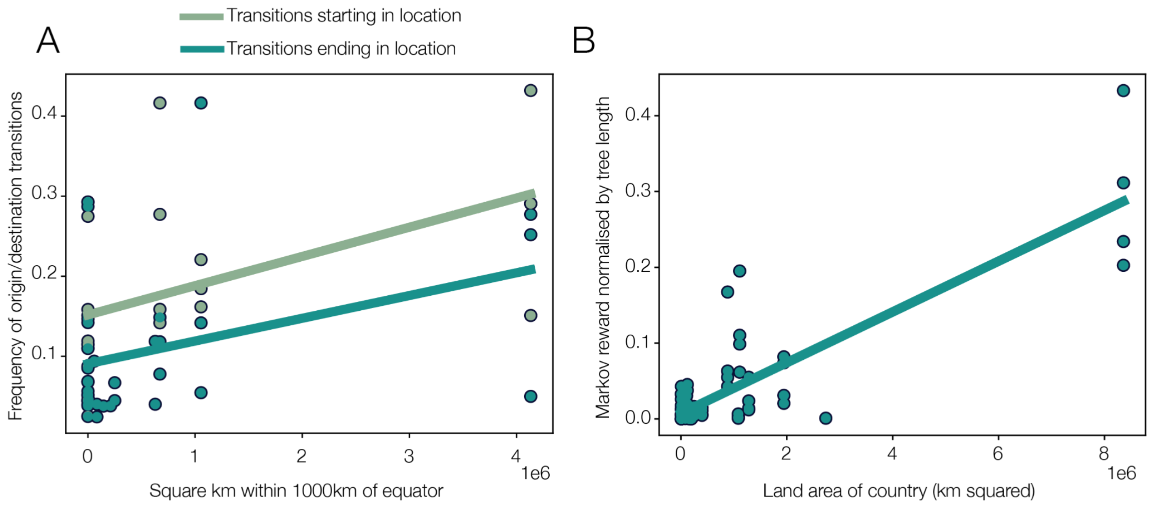


***Figure S8**: **Correlations for Markov jumps and rewards**. (**A**) Linear regression of the frequency of significant transitions that start (green) and end (blue) across the posterior against the square kilometres within 1000km of the equator for each country, using an equal area map projection; (**B**) Markov rewards against the land area of each country in km$^2$ using an equal area map projection.*

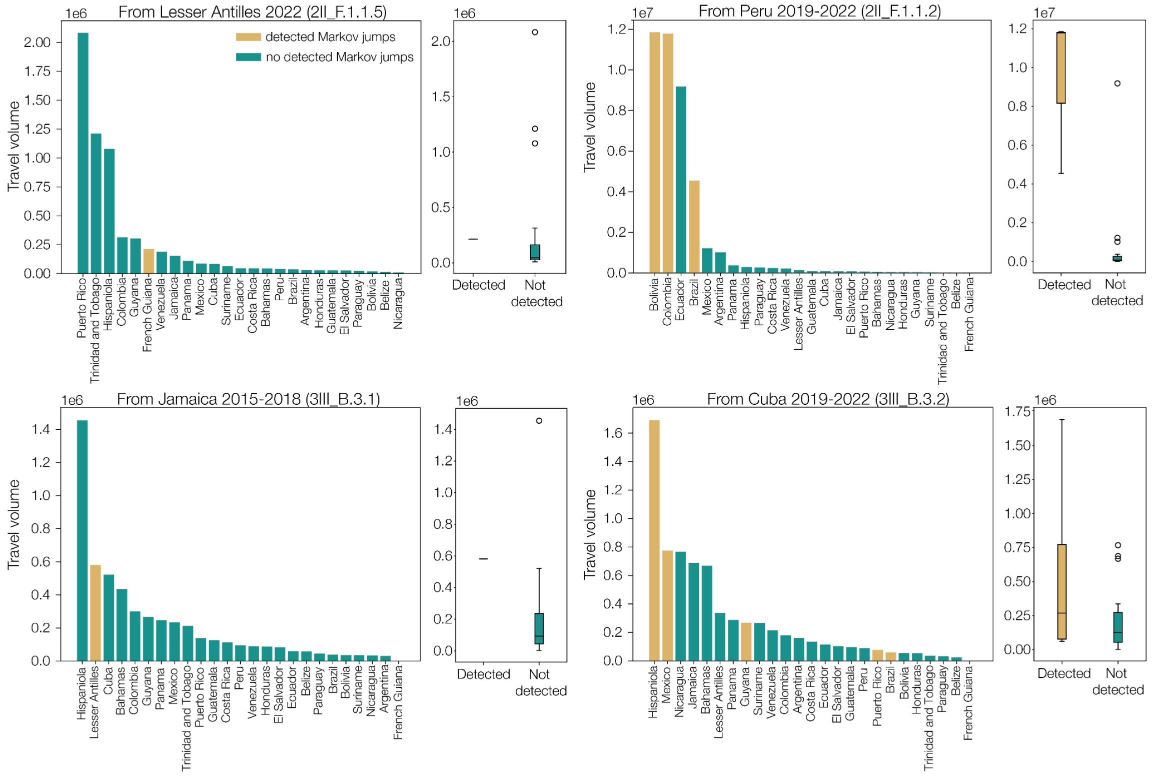


***Figure S9: Travel volume for smaller introductions** Bars show travel volume from inferred origin location of introduction for relevant time period. Blue bars are destinations with no detected Markov jumps in the introduction, yellow are destinations with detected Markov*

*jumps. Box plots show spread of travel volume, split by destinations with and without jumps detected.*

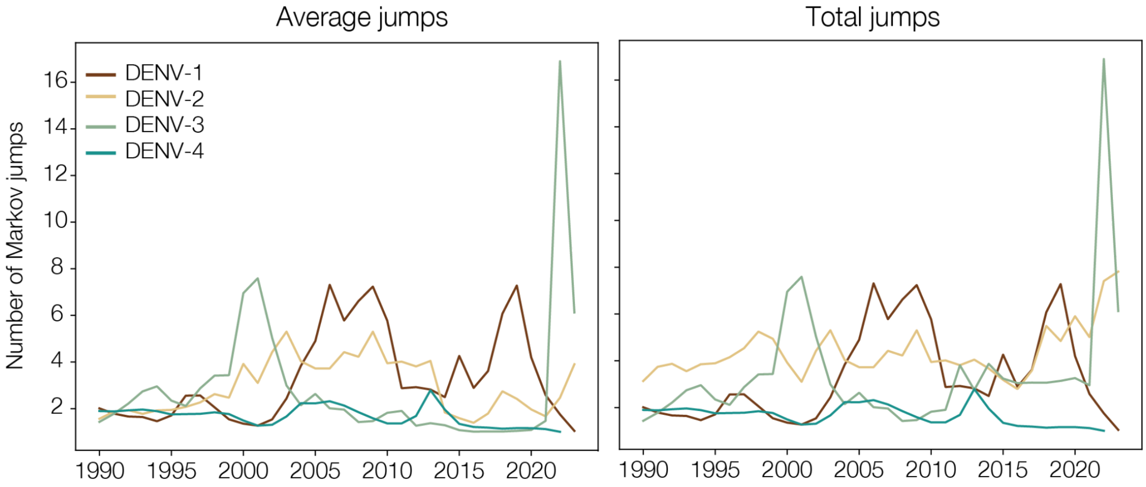


***Figure S10*** *Number of Markov jumps across all introductions into the Americas by serotype, average on the left and total on the right.*
